# LPMOcat: an interactive database for comparative structural analysis of lytic polysaccharide monooxygenases

**DOI:** 10.64898/2026.11.12.766080

**Authors:** Katarzyna Szleper-Shendy, Carlos Sequeiros-Borja, Patryk Klytta, Artur Góra

## Abstract

**Summary:** LPMOcat is a comprehensive resource of 161 characterised lytic polysaccharide monooxygenases (LPMOs) from all eight CAZy families, in which an automated pipeline integrates data from public databases and, based on a structural alignment, annotates common structural features of LPMOs (e.g. histidine brace, second-sphere residues, loop regions), and calculates per-residue properties (e.g. flexibility, hydrophobicity, hole-hopping pathways). The web application summarises all the gathered information in plots integrated with a similarity tree and allows users to filter the catalogue and compare any set of LPMOs to investigate their structure– function relationships.

**Availability and implementation:** LPMOcat is freely available at https://lpmo-cat.polsl.pl. The source code is available at https://github.com/TUNNELING-GROUP/LPMOcat and archived at Zenodo https://doi.org/10.5281/zenodo.23698332.

**Supplementary information:** Supplementary data available at the journal’s web site.

## Introduction

Lytic polysaccharide monooxygenases (LPMOs) were the first enzymes found to oxidatively cleave glycosidic bonds in polysaccharides, such as cellulose and chitin (Vaaje-Kolstad *et al*. 2010, Quinlan *et al*. 2011, Phillips *et al*. 2011). They are key players in biomass degradation (Chylenski *et al*. 2019), although their biological roles extend beyond it (Vandhana *et al*. 2022). In the Carbohydrate-Active enZymes (CAZy) database, LPMOs are classified based on sequence similarity into eight families of its Auxiliary Activities class (AA9–AA11 and AA13–AA17) (Levasseur *et al*. 2013, Drula *et al*. 2022). LPMOs appear mainly in fungi and bacteria, but also in oomycetes, animals, plants and viruses, and the families differ in their taxonomic distribution. All LPMOs share an immunoglobulin-like β-sandwich core and a surface-exposed active site, in which a copper ion is coordinated by two histidines, the so-called histidine brace (Figure 1A). Much of the structural diversity between and within families lies in the loops connecting the strands of the core, which shape the flat substrate-binding surface and contain many of the second-sphere residues surrounding the histidine brace. These differences are thought to tune LPMO reactivity and determine substrate specificity (e.g. cellulose, chitin, hemicelluloses or starch) and regioselectivity (oxidation at C1, C4 or both) (Vaaje-Kolstad *et al*. 2017, Munzone *et al*. 2024). The activity and stability of LPMOs are further modulated by hole-hopping pathways. These pathways, consisting mainly of tyrosine and tryptophan residues, connect the surface-exposed active site, *via* buried residues in the core, with distant sites on the protein surface, and can transfer electrons or holes formed during redox reactions, either supporting catalysis or protecting the enzyme from oxidative damage (Bissaro *et al*. 2017) (Ayuso-Fernández *et al*. 2024, 2026, Sayler *et al*. 2025).

**Figure 1.**
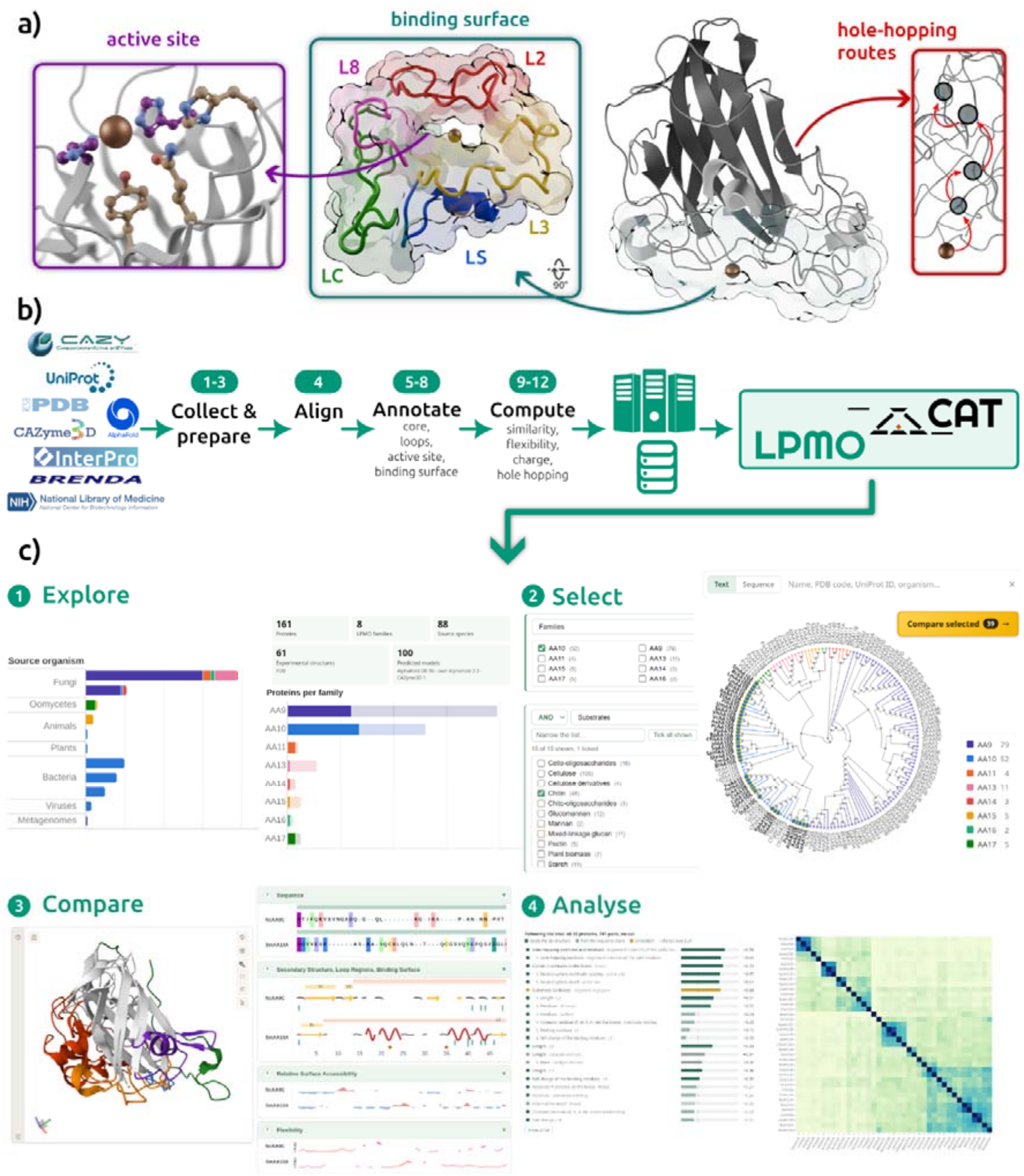
LPMOcat: structural features, data pipeline and web application. A) Structural features annotated uniformly for every LPMO in the catalogue, shown on NcAA9C (4D7U): the copper active site with the histidine brace and second-sphere residues; the substrate-binding surface with the loop regions L2, L3, LS, L8 and LC (view rotated by 90°); and predicted hole-hopping routes from the copper. B) An automated 12-step pipeline collects characterised LPMOs and their data from public resources, prepares one structure per protein, superposes all structures in a single structural alignment, annotates structural elements and calculates per-residue properties. The results are stored in a database queried by the LPMOcat web application. C) Example of the results of the workflow in the web application.

Since their discovery, LPMOs have attracted wide interest and have been characterised functionally (Eijsink *et al*. 2019), structurally (Frandsen and Lo Leggio 2016) and computationally (Hagemann and Hedegård 2023). The resulting data are curated in several public databases. In CAZy (Drula *et al*. 2022) tens of thousands of LPMO sequences are currently available, the PDB (Burley *et al*. 2023) contains experimental structures for about 60 of them and UniProt (The UniProt Consortium 2025) and BRENDA (Hauenstein *et al*. 2026) collect their functional annotations. However, these databases are separate, and to our knowledge there is no resource available that integrates structural and functional information on LPMOs in a way that allows them to be explored and compared across families. Moreover, although the loop regions (L2, L3, LS, L8 and LC, Figure 1A) are used to describe LPMOs across families (Zhou and Zhu 2020), their boundaries are assigned individually for each structure rather than by a common definition, and second-sphere motifs have been described only for individual LPMOs, mainly for AA9 and AA10 LPMOs (Hall *et al*. 2023, 2024). Similarly, residues that contact the substrate have been determined experimentally for only a few LPMOs, mostly from AA9 and AA10 (Aachmann *et al*. 2012, Frandsen *et al*. 2016), so the binding residues of most LPMOs remain undefined.

To facilitate visualisation and comparison of LPMO properties and structural features, we developed LPMOcat (https://lpmo-cat.polsl.pl), an interactive web resource, integrating functional annotations with structurally aligned experimental structures and predicted models. The current release contains 161 proteins from all known families, selected based on experimental evidence (biochemical characterization or a solved structure) reported in CAZy, UniProtKB/Swiss-Prot, BRENDA or the literature. An automated pipeline (Figure 1B) collects data from public databases, selects one 3D structure per protein and superposes all structures in a single alignment. Based on it, common structural properties of LPMOs, such as core strands, loop regions, histidine brace, second-sphere residue motifs and substrate-binding residues are uniformly annotated. The pipeline also calculates per-residue properties (flexibility, solvent accessibility, charge and hydrophobicity) and predicts hole-hopping pathways. Additionally, the pairwise similarity between all proteins is computed. The user-friendly web application presents these data interactively, allowing users to explore the whole catalogue, as well as to select any set of LPMOs for detailed structural comparison (Figure 1C).

## Methods and implementation

LPMOcat contains characterised LPMOs only, i.e. proteins with experimental evidence of activity or an experimentally solved structure. It is built by an automated pipeline of 12 steps, whose output is loaded into a database, which is further queried and displayed by the web application (Figure 1B). First, these proteins are collected from CAZy (Drula *et al*. 2022), reviewed entries in UniProtKB/Swiss-Prot (The UniProt Consortium 2025), BRENDA (Hauenstein *et al*. 2026) and in-house database. For each protein, accessions, InterPro domain annotations (Blum *et al*. 2025), EC numbers, NCBI taxonomy (Schoch *et al*. 2020) and reported substrates and regioselectivity are retrieved, each kept with its source publication. Each protein is represented by one structure, an experimental structure from the PDB (Burley *et al*. 2023) when available, or a predicted model, mostly from AlphaFold DB (Varadi *et al*. 2022, 2024). All structures are prepared uniformly. Each is reduced to a single chain with standard residues and trimmed to start at the copper-coordinating N-terminal histidine, and short unresolved loops are rebuilt with MODELLER (Šali and Blundell 1993). They are then superposed in a multiple structural alignment with US-align (Zhang C *et al*. 2022). Based on the alignment, the same structural elements are defined in every protein across all families: the end of the catalytic domain, disulfide bonds, the β-sandwich core, the substrate-binding loops L2, L3, LS, L8 and LC, the histidine brace with second-sphere residues and geometrically approximated substrate-binding surface. In addition, each residue is annotated with five properties: secondary structure using mkdssp (Kabsch and Sander 1983, Hekkelman *et al*. 2025); relative solvent accessibility using Biopython (Shrake and Rupley 1973, Cock *et al*. 2009); flexibility using CABS-flex (Kurcinski *et al*. 2019), ProDy (Zhang S *et al*. 2021) and B-factors or pLDDT values (Jumper *et al*. 2021); partial charge using PDB2PQR (Dolinsky *et al*. 2004); and participation in predicted hole-hopping pathways using EHPath (Teo *et al*. 2019). Finally, pairwise distances between catalytic domains are computed with Biopython (Cock *et al*. 2009) from BLOSUM62 scores (Henikoff and Henikoff 1992) over the aligned positions, and are used to build a UPGMA similarity tree used for navigation and selection.

The web application is built with Django framework and uses SQLite database to store protein records. In the web application, structures are displayed with Mol* (Sehnal *et al*. 2021), plots are rendered using Plotly (Plotly Technologies Inc 2015) and the similarity tree with D3.js (Bostock, Ogievetsky, and Heer 2011). A detailed description of the pipeline steps and the web application is provided in the Supplementary Information (sections “Pipeline description” and “Database and web application”) and in the pipeline documentation (https://lpmo-cat.polsl.pl/docs/).

## Usage and results

### Catalogue content

The current release contains 161 LPMOs from all eight families, represented by 61 experimental structures and 100 predicted models (Supplementary Table S1). Most come from fungi and bacteria (Supplementary Table S2), and 142 have activity data, mainly on cellulose and chitin, with substrates and regioselectivity reported in papers for 82 of them and otherwise derived from EC numbers (Supplementary Tables S3– S4). The consistent annotation reveals family-specific patterns around the copper (Supplementary Tables S5–S7). These and further properties of the catalytic domains, such as length, hydrophobicity, charge and loop lengths (Supplementary Figure S1), can be explored interactively in the Catalogue overview (https://lpmo-cat.polsl.pl/catalogue/, Figure 1C, step 1), where each chart can be split by family or restricted to a set of residues, and linked to the similarity tree.

### Selection of LPMOs for comparison

Proteins can be selected for comparison in several ways: from the charts of the overview, from nodes or branches of the similarity tree, by searching for a name, organism, accession, sequence or motif, or by combining filters for annotations (e.g. family, substrate, regioselectivity, motifs or loop lengths) as criteria with boolean operators AND, OR and NOT (Figure 1C, step 2). Up to 50 proteins, or a whole family, can be compared at once. Selected proteins can also be saved as up to three named groups, which the comparison page marks in the table, the plots and the 3D viewer.

### Comparative analysis and visualization

The comparison page shows the selected proteins together (Figure 1C, step 3). A sortable table, whose columns can be shown or hidden, lists their annotations and computed properties of the catalytic domain (e.g. net charge, isoelectric point, hydrophobicity, disulfide bonds), with substrates and regioselectivity linked to the publications that report them, including substrates tested without activity (Supplementary Figure S2). Below the table, all structures are superposed in a 3D viewer and can be coloured by any of the per-residue properties, with the histidine brace, second-sphere residues, binding surface, loops, disulfide bonds and hole-hopping paths shown on demand (Supplementary Figure S3). The per-residue data are drawn as plots along the alignment, one row per protein, so that equivalent positions lie in one column across families. The plots and the viewer are linked. Pointing at a residue in a plot highlights it in the structures and vice versa, and each plot can colour the structures (Supplementary Figure S3).

Finally, on the comparison page of LPMOcat, the selection overview allows the compared proteins to be analysed further, either on their own or against the rest of the catalogue (Figure 1C, step 4, Supplementary Figure S4). This overview is built around the similarity tree, pruned to the compared proteins and annotated with their structure source, regioselectivity, second-sphere residues and brace motifs. Proteins or branches picked in the tree, or groups defined by the user or by cutting the tree, can be characterised by 75 features (Supplementary Table S8), such as the size, charge and hydrophobicity of the catalytic domain and its regions (surface, loops, substrate-binding residues), loop flexibility, second-sphere and brace motifs, hole-hopping positions, reported substrates and taxonomy, and others. The features are ranked by how closely they follow the tree, how well they separate the groups, or how well they distinguish the picked proteins from the rest of the comparison, of their families or of the catalogue, each against a permutation-based threshold. The rankings show which features proteins share, rather than causal relationships or functional. The tree is linked to the overview charts and to two views along the structural alignment: a sequence logo (Schneider and Stephens 1990) of the catalytic domain, annotated with the brace, second-sphere residues, binding surface and loops, and a heatmap of the pairwise similarity of the proteins for a chosen property and region, e.g. charge of the loops or flexibility of the hole-hopping residues, ordered by the tree or clustered by the property itself. Pointing at a bar, a letter of the logo or a cell of the heatmap highlights the corresponding proteins in the tree.

## Conclusions

LPMOcat is an interactive web resource that collects characterised LPMOs from all families and allows them to be visualised and compared. Based on a single structural alignment, it uniformly annotates common structural elements of LPMOs, such as the core strands, loop regions, histidine brace, second-sphere residues and substrate-binding surface. Together with per-residue properties, such as flexibility, solvent accessibility, charge and hole-hopping pathways, these annotations can be compared position by position across families for any selection of proteins and related to their reported substrates and regioselectivity. LPMOcat can therefore support the design of experiments, the interpretation of results and the formulation of hypotheses on structure-function relationships. As the pipeline is openly available and documented step by step, the catalogue can be updated as new LPMOs are characterised, and the approach can be reused for other enzyme families.

## Funding

The work of K.S.S. and A.G. was supported by NewCat project under the European Commission EIC Pathfinder program (Grant No. 101046815), work of C.S.B and P.K. by National Scientific Centre, Poland, grant number UMO-2024/55/B/ST4/02371. K.S.S. work was additionally supported by funding from the NAWA STER Program, within the framework of the NIH4DS - New International Horizons for Doctoral School project. We gratefully acknowledge Polish high-performance computing infrastructure PLGrid (HPC Centers: ACK Cyfronet AGH) for providing computer facilities and support within computational grants no. PLG/2022/015696, PLG/2024/017171 and PLG/2025/018297.

We thank Michał Chyski and Wiktor Stolarz for developing the first prototypes of the LPMOcat website, and Andrea K. Hönikl for curating the LPMOs added to the catalogue from the literature. We also thank Dr. Gamze Tanriver for insightful discussions, valuable feedback and assistance with testing the prototype version.

Claude (Anthropic) was used to assist with editing the manuscript text and with writing the code. The authors reviewed, tested and revised all text and code and take full responsibility for the content of this article and the software.

## Notes

https://doi.org/10.5281/zenodo.23698332

https://github.com/TUNNELING-GROUP/LPMOcat

https://lpmo-cat.polsl.pl

### Competing Interest Statement

The authors have declared no competing interest.

## References

Aachmann FL, Sørlie M, Skjåk-Bræk G et al. NMR structure of a lytic polysaccharide monooxygenase provides insight into copper binding, protein dynamics, and substrate interactions. Proc Natl Acad Sci U S A 2012;109(46):18779–84. 10.1073/pnas.1208822109.

Ayuso-Fernández I, Emrich-Mills TZ, Golten O et al. Redox robustness drives LPMO evolution. Proc Natl Acad Sci 2026;123(3):e2521617123. 10.1073/pnas.2521617123.

Ayuso-Fernández I, Emrich-Mills TZ, Haak J et al. Mutational dissection of a hole hopping route in a lytic polysaccharide monooxygenase (LPMO). Nat Commun 2024;15(1). 10.1038/s41467-024-48245-w.

Bissaro B, Røhr ÅK, Müller G et al. Oxidative cleavage of polysaccharides by monocopper enzymes depends on H2O2. Nat Chem Biol 2017;13(10):1123–8. 10.1038/nchembio.2470.

Blum M, Andreeva A, Florentino LC et al. InterPro: the protein sequence classification resource in 2025. Nucleic Acids Res 2025;53(D1):D444–56. 10.1093/nar/gkae1082.

Bostock M, Ogievetsky V, Heer J. D3 Data-Driven Documents. IEEE Trans Vis Comput Graph 2011;17(12):2301–9. 10.1109/TVCG.2011.185.

Burley SK, Bhikadiya C, Bi C et al. RCSB Protein Data Bank (RCSB.org): delivery of experimentally-determined PDB structures alongside one million computed structure models of proteins from artificial intelligence/machine learning. Nucleic Acids Res 2023;51(D1):D488–508. 10.1093/nar/gkac1077.

Chylenski P, Bissaro B, Sørlie M et al. Lytic Polysaccharide Monooxygenases in Enzymatic Processing of Lignocellulosic Biomass. ACS Catal 2019;9(6):4970–91. 10.1021/acscatal.9b00246.

Cock PJA, Antao T, Chang JT et al. Biopython: freely available Python tools for computational molecular biology and bioinformatics. Bioinformatics 2009;25(11):1422–3. 10.1093/bioinformatics/btp163.

Dolinsky TJ, Nielsen JE, McCammon JA et al. PDB2PQR: an automated pipeline for the setup of Poisson-Boltzmann electrostatics calculations. Nucleic Acids Res 2004;32(Web Server):W665–7. 10.1093/nar/gkh381.

Drula E, Garron ML, Dogan S et al. The carbohydrate-active enzyme database: Functions and literature. Nucleic Acids Res 2022;50(D1):D571–7. 10.1093/nar/gkab1045.

Eijsink VGH, Petrovic D, Forsberg Z et al. On the functional characterization of lytic polysaccharide monooxygenases (LPMOs). Biotechnol Biofuels 2019;12(1). 10.1186/s13068-019-1392-0.

Frandsen KEH, Lo Leggio L. Lytic polysaccharide monooxygenases: A crystallographer’s view on a new class of biomass-degrading enzymes. IUCrJ 2016;3:448–67. 10.1107/S2052252516014147.

Frandsen KEH, Simmons TJ, Dupree P et al. The molecular basis of polysaccharide cleavage by lytic polysaccharide monooxygenases. Nat Chem Biol 2016;12(4):298–303. 10.1038/nchembio.2029.

Hagemann MM, Hedegård ED. Molecular Mechanism of Substrate Oxidation in Lytic Polysaccharide Monooxygenases: Insight from Theoretical Investigations. Chem - Eur J 2023;29(7). 10.1002/chem.202202379.

Hall KR, Joseph C, Ayuso-Fernández I et al. A Conserved Second Sphere Residue Tunes Copper Site Reactivity in Lytic Polysaccharide Monooxygenases. J Am Chem Soc 2023;145(34):18888–903. 10.1021/jacs.3c05342.

Hall KR, Mollatt M, Forsberg Z et al. Impact of the Copper Second Coordination Sphere on Catalytic Performance and Substrate Specificity of a Bacterial Lytic Polysaccharide Monooxygenase. ACS Omega 2024;9(21):23040–52. 10.1021/acsomega.4c02666.

Hauenstein J, Jeske L, Jäde A et al. BRENDA in 2026: a Global Core Biodata Resource for functional enzyme and metabolic data within the DSMZ Digital Diversity. Nucleic Acids Res 2026;54(D1):D527–34. 10.1093/nar/gkaf1113.

Hekkelman ML, Salmoral DÁ, Perrakis A et al. DSSP 4: FAIR annotation of protein secondary structure. Protein Sci 2025;34(8):e70208. 10.1002/pro.70208.

Henikoff S, Henikoff JG. Amino acid substitution matrices from protein blocks. Proc Natl Acad Sci 1992;89(22):10915–9. 10.1073/pnas.89.22.10915.

Jumper J, Evans R, Pritzel A et al. Highly accurate protein structure prediction with AlphaFold. Nature 2021;596(7873):583–9. 10.1038/s41586-021-03819-2.

Kabsch W, Sander C. Dictionary of protein secondary structure: Pattern recognition of hydrogen-bonded and geometrical features. Biopolymers 1983;22(12):2577–637. 10.1002/bip.360221211.

Kurcinski M, Oleniecki T, Ciemny MP et al. CABS-flex standalone: a simulation environment for fast modeling of protein flexibility. Bioinformatics 2019;35(4):694–5. 10.1093/bioinformatics/bty685.

Levasseur A, Drula E, Lombard V et al. Expansion of the enzymatic repertoire of the CAZy database to integrate auxiliary redox enzymes. Biotechnol Biofuels 2013;6(1):41. 10.1186/1754-6834-6-41.

Munzone A, Eijsink VGH, Berrin JG et al. Expanding the catalytic landscape of metalloenzymes with lytic polysaccharide monooxygenases. Nat Rev Chem 2024;8(2):106–19. 10.1038/s41570-023-00565-z.

Phillips CM, Beeson WT, Cate JH et al. Cellobiose Dehydrogenase and a Copper-Dependent Polysaccharide Monooxygenase Potentiate Cellulose Degradation by Neurospora crassa. ACS Chem Biol 2011;6(12):1399–406. 10.1021/cb200351y.

Plotly Technologies Inc. Collaborative data science. Montréal, QC, 2015. https://plot.ly.

Quinlan RJ, Sweeney MD, Lo Leggio L et al. Insights into the oxidative degradation of cellulose by a copper metalloenzyme that exploits biomass components. Proc Natl Acad Sci 2011;108(37):15079–84. 10.1073/pnas.1105776108.

Šali A, Blundell TL. Comparative Protein Modelling by Satisfaction of Spatial Restraints. J Mol Biol 1993;234(3):779–815. 10.1006/jmbi.1993.1626.

Sayler RI, Thomas WC, Rose AJ et al. Electron transfer in polysaccharide monooxygenase catalysis. Proc Natl Acad Sci U S A 2025;122(1). 10.1073/pnas.2411229121.

Schneider TD, Stephens RM. Sequence logos: a new way to display consensus sequences. Nucleic Acids Res 1990;18(20):6097–100. 10.1093/nar/18.20.6097.

Schoch CL, Ciufo S, Domrachev M et al. NCBI Taxonomy: a comprehensive update on curation, resources and tools. Database 2020;2020:baaa062. 10.1093/database/baaa062.

Sehnal D, Bittrich S, Deshpande M et al. Mol* Viewer: modern web app for 3D visualization and analysis of large biomolecular structures. Nucleic Acids Res 2021;49(W1):W431–7. 10.1093/nar/gkab314.

Shrake A, Rupley JA. Environment and exposure to solvent of protein atoms. Lysozyme and insulin. J Mol Biol 1973;79(2):351–71. 10.1016/0022-2836(73)90011-9.

Teo RD, Wang R, Smithwick ER et al. Mapping hole hopping escape routes in proteins. Proc Natl Acad Sci U S A 2019;116(32):15811–6. 10.1073/pnas.1906394116.

The UniProt Consortium. UniProt: the Universal Protein Knowledgebase in 2025. Nucleic Acids Res 2025;53(D1):D609–17. 10.1093/nar/gkae1010.

Vaaje-Kolstad G, Forsberg Z, Loose JS et al. Structural diversity of lytic polysaccharide monooxygenases. Curr Opin Struct Biol 2017;44:67–76. 10.1016/j.sbi.2016.12.012.

Vaaje-Kolstad G, Westereng B, Horn SJ et al. An oxidative enzyme boosting the enzymatic conversion of recalcitrant polysaccharides. Science 2010;330(6001):219–22. 10.1126/science.1192231.

Vandhana TM, Reyre JL, Sushmaa D et al. On the expansion of biological functions of lytic polysaccharide monooxygenases. New Phytol 2022;233(6):2380–96. 10.1111/nph.17921.

Varadi M, Anyango S, Deshpande M et al. AlphaFold Protein Structure Database: massively expanding the structural coverage of protein-sequence space with high-accuracy models. Nucleic Acids Res 2022;50(D1):D439–44. 10.1093/nar/gkab1061.

Varadi M, Bertoni D, Magana P et al. AlphaFold Protein Structure Database in 2024: providing structure coverage for over 214 million protein sequences. Nucleic Acids Res 2024;52(D1):D368–75. 10.1093/nar/gkad1011.

Zhang C, Shine M, Pyle AM et al. US-align: universal structure alignments of proteins, nucleic acids, and macromolecular complexes. Nat Methods 2022;19(9):1109–15. 10.1038/s41592-022-01585-1.

Zhang S, Krieger JM, Zhang Y et al. ProDy 2.0: increased scale and scope after 10 years of protein dynamics modelling with Python. Bioinformatics 2021;37(20):3657–9. 10.1093/bioinformatics/btab187.

Zhou X, Zhu H. Current understanding of substrate specificity and regioselectivity of LPMOs. Bioresour Bioprocess 2020;7(1). 10.1186/s40643-020-0300-6.

