## Supplementary data for "LPMOcat: an interactive database for comparative structural analysis of lytic polysaccharide monooxygenases"

### Pipeline description

#### Protein retrieval (step 1)

The core list of proteins in LPMOcat is built from the eight LPMO families within the CAZy database (AA9–AA11, AA13–AA17) (Levasseur *et al.* 2013, Drula *et al.* 2022). For each family, CAZy marks the members whose activity has been demonstrated experimentally on its "characterized" page, and those with a structure in the PDB on its "structure" page, and those two pages were used as source. For each entry, the protein name, family, source organism, EC numbers, GenBank and UniProt accessions, PDB codes and the characterisation papers CAZy cites are retrieved. On 21 October 2026, CAZy listed 134 entries: 121 characterised, of which 49 also had a structure, and 13 with a structure only.

Characterised LPMOs not yet listed in CAZy are added from three further sources. From UniProt (The UniProt Consortium 2025), we take reviewed (Swiss-Prot) entries with an LPMO EC number (EC 1.14.99.53–56) whose activity is supported by experimental evidence (ECO:0000269) and a cited publication: 73 of 114 matching entries in release 2026_03, of which 28 were not in CAZy. From BRENDA (Hauenstein *et al.* 2026), we take proteins with these EC numbers that have a UniProt accession and a substrate record citing a paper on that organism (2 of 56 proteins in release 2026.1 were not already included). Finally, our in-house database adds two unreviewed UniProt entries (CtPMO1, CgAA15A) and three proteins without a public model (BaAA9A, PchAA9E, ScAA9A; see step 2), giving 169 entries

#### Data fetching, structure selection and preparation (steps 2-3)

Each protein is represented by one 3D structure (step 2). An experimental structure from the Protein Data Bank (Burley *et al.* 2023) is preferred; among its entries, wild-type X-ray structures with copper in the active site are chosen when their resolution is within 1 Å of the best available, and further ties are broken by resolution, completeness and chain length. Otherwise, the AlphaFold DB model (Varadi *et al.* 2024) of the protein’s UniProt entry, or of a UniProt entry with an identical sequence (UniParc), is used, and when none exists, a CAZyme3D model (Siva Shanmugam and Yin 2025). Three proteins without a suitable public model (BaAA9A and PchAA9E available only in MycoCosm, and ScAA9A, whose UniProt sequence derives from an outdated gene model) were modelled with AlphaFold 3 (Abramson *et al.* 2024). Entries with identical sequences are kept once (3 removed), strain variants (≥ 95 % identity, same species) are merged into the protein with a structure (3), and entries without any structure are excluded (2). The current release contains 161 proteins: 61 experimental structures, 96 AlphaFold DB models, 1 CAZyme3D model and 3 AlphaFold 3 models.

For each protein, all its UniProt accessions (including identical-sequence entries and merged strain variants), domain annotations from UniProt and InterPro (Blum *et al.* 2025), EC numbers and the NCBI Taxonomy lineage (Schoch *et al.* 2020) are then collected. Substrates and oxidised positions reported by CAZy, Swiss-Prot, BRENDA and the literature are stored as separate claims, each with its source and supporting paper; negative results are kept, and disagreements between sources are flagged rather than resolved. In the current release, 142 of the 161 proteins have activity data, supported by 160 publications.

All structures are prepared in the same way (step 3). One model, one chain and one alternate location are kept, modified residues are replaced by their parent amino acids and recorded (e.g. the methylated N-His of fungal LPMOs, present in 10 structures), and only the protein and its active-site metal are retained. Internal gaps of up to 30 residues are rebuilt with MODELLER (Šali and Blundell 1993) (5 of the 61 experimental structures). Because the free amino group of the N-terminal histidine (N-His) coordinates the copper, every structure is trimmed to start at this residue, using the annotated signal peptide or a SignalP 6.0 prediction (Teufel *et al.* 2022). C-terminal expression tags are removed (6 structures).

#### Structural alignment (step 4)

All prepared structures are aligned with US-align (version 20241108) in multiple-structure mode (-mm 4) (Zhang C *et al.* 2022), which assigns every residue to a column of one alignment and superposes all structures in one coordinate frame. Because LPMOs of different families share only 2-20 % sequence identity, the alignment is not refined by sequence. It has an average pairwise TM-score of 0.56 and RMSD of 3.1 Å over 148 matched residues per pair; of its 710 columns within the catalytic domains, 41 are occupied in all 161 proteins.

#### Annotation of structural elements (steps 5-8)

The structures are annotated so that the same name denotes the equivalent structural element in every protein, across all eight families. First, secondary structure is assigned with DSSP (mkdssp 4.4) (Kabsch and Sander 1983, Hekkelman *et al.* 2025) and solvent accessibility is computed with the Shrake-Rupley algorithm in Biopython (Shrake and Rupley 1973, Cock *et al.* 2009); residues with a relative solvent accessibility (RSA) (Tien *et al.* 2013) of at least 25 % are counted as surface exposed (step 5).

Next, the end of the catalytic domain, the disulfide bonds and the histidine brace are identified (step 6). The catalytic-domain end is taken from the UniProt and InterPro annotation for AlphaFold models and is otherwise placed at the family consensus (the median alignment column of the annotated ends); it is then shifted out of long coil segments and never placed between the two cysteines of a disulfide bond. The two brace histidines, the N-His and the internal His, are identified from the metal in the 57 structures that contain one, and elsewhere at the corresponding alignment columns. In structures without a metal, the copper position is approximated by a pseudo-copper at the centroid of the five brace nitrogens. The brace is conserved across the alignment, with a few exceptions (e.g. LsAA9B, which has an arginine in place of the N-His).

The conserved fold is then described (step 7). The β-sandwich core is defined from the alignment columns where a β-strand is common across families, weighted so that large families do not dominate, and checked by strand pairing within the sheets. Core strands are numbered β1–β9, with β2 being the hairpin partner of β1 found only in AA9, so that the same number marks the equivalent strand in every family. The loops described in the literature (e.g. Zhou and Zhu 2020) are defined as the segments between core strands: L2 between β2 (or β1) and β3, L3 between β4 and β5, LS between β6 and β7, L8 between β8 and β9, and LC after β9. Because LPMOs act on mostly flat crystalline polysaccharides, the substrate-binding surface is approximated by the plane, among many laid against the copper face, that touches the most residues around the copper with the copper at most 3 Å below it; its solvent-accessible contact residues form the binding surface.

Finally, the second-sphere residues around the copper are annotated (step 8): the aromatic residue at the axial position of the copper (e.g. Tyr in AA9, mostly Phe in AA10) (Frandsen and Lo Leggio 2016); side chains in hydrogen-bond, π-stacking or cation–π contact with the brace (within 3.5 Å); the second-sphere hydrogen-bonding motif (e.g. H-X8-Q-X-Y in AA9, E...N-X-F and R...E-X-F in AA10) (Span and Marletta 2015, Span *et al.* 2017, Laurent *et al.* 2019, Hall *et al.* 2023, 2024, Batka *et al.* 2024); the residue shielding the solvent-facing axial position of the copper (Ala in most AA10s) (Hemsworth, Davies, and Walton 2013); and the sequence motifs following the two brace histidines, such as H-X-G-P after the internal His (Laurent *et al.* 2019) and H-G-X4-P-X2-R after the N-His, described here.

#### Sequence similarity (step 9)

Pairwise distances between the catalytic domains are computed from the structural alignment with Biopython (Cock *et al.* 2009): over the columns where both proteins have a residue, the BLOSUM62 score of the aligned pairs is divided by the higher of the two self-scores and subtracted from 1. From these distances, the web application builds a tree by average linkage (UPGMA) (Sokal and Michener 1958). The tree groups proteins for navigation and comparison but is not a phylogeny; at 2-20 % identity between families, their branching order is not resolved.

#### Per-residue properties (steps 10–12)

Flexibility is estimated with two methods, shown separately because their scales differ (step 10): mean-square fluctuations from a Gaussian network model of the Cα atoms using ProDy (Zhang S *et al.* 2021) (10 Å cutoff, 20 slowest modes), and root-mean-square fluctuations from coarse-grained CABS-flex simulations with default settings (Kurcinski *et al.* 2019). In addition, the Cα B-factors of experimental structures and the per-residue confidence (pLDDT) of

AlphaFold models (Jumper *et al.* 2021) are provided as indicators of local disorder and model reliability.

Atomic partial charges are assigned with PDB2PQR (version 2.1.1, AMBER force field) (Dolinsky *et al.* 2004) (step 11). All proteins are treated in the same metal-free state, since most structures are models without copper, and residues take standard protonation states without pKa prediction. The two brace histidines are protonated so that their copper-coordinating ring nitrogens remain free. The N-terminal amine is protonated (–$NH3^{+}$) as in a free N-terminus, although in the copper-bound enzyme it coordinates the copper as a neutral amine. This simplification avoids parametrising the copper site and adds the same +1 to the net charge of every protein, so comparisons between proteins are not affected. The charge of a residue, or of a group of residues such as a loop, is the sum of its atomic charges.

Residues that may transfer electrons (holes) between the protein surface and the copper are predicted with EHPath (Teo *et al.* 2019), as in a recent LPMO study (Sayler *et al.* 2025) (step 12). Surface-exposed (RSA ≥ 10 %) Tyr, Trp and His residues act as donors; buried Trp, and buried Tyr whose hydroxyl lies within 3.5 Å of the copper or 4 Å of a proton acceptor, act as bridging residues. Paths of at most six nodes, including donor and copper, are searched from each donor, the five best paths per donor are kept, and all paths are ranked by mean residence time. For the copper and histidine, which EHPath does not parametrise, λ = 1.0 eV and an oxidation potential of 1.0 V are assumed, so the ranking serves to compare pathways rather than to estimate absolute rates.

### Database and web application

The pipeline output is loaded into a relational database (SQLite, managed by Django 5.1; Django Software Foundation 2024), with one record per protein holding its annotations, activity claims and references, and linked per-residue records for surface accessibility, flexibility, charge and hole-hopping paths. During loading, the substrates and regioselectivity of each protein are taken from the claims reported in papers or, where none exist, derived from its EC numbers using the IUBMB definitions (ExplorEnz) (McDonald, Boyce, and Tipton 2009). References from the structure entries, CAZy and UniProt are merged and deduplicated by DOI and PubMed ID, and the domains annotated in UniProt, including carbohydrate-binding modules, are mapped onto the structure residues via SIFTS (Dana *et al.* 2019) or by sequence alignment.

The web application uses Django on the server and JavaScript in the browser. Structures are displayed with Mol* (Sehnal *et al.* 2021) through the PDBe Molstar plugin, plots with Plotly (Plotly Technologies Inc 2015) and the similarity tree with D3.js (Bostock, Ogievetsky, and Heer 2011), built in the browser from the distance matrix of step 9. Comparisons of any selection are computed on request, including the pairwise similarity of the proteins for one property over one region (NumPy, SciPy) and a sequence logo of every alignment column.

The compared proteins are described by 75 features (Table S8). Features are ranked in three ways: by how closely differences in the feature follow distances in the similarity tree (Mantel test with Spearman correlation); by how well they separate the groups formed by cutting the tree ($\eta^2$ for numerical, Cramér's V for categorical features); and by how well they discriminate the selected proteins from a reference set ($|2 \cdot AUC - 1|$ for numerical features, total variation distance for categorical ones). Hole-hopping positions, which are sets of alignment columns, are compared by Jaccard distance (PERMANOVA $R^2$ for groups). For each ranking, a permutation-based threshold is the 95th percentile of the best score over all features in 1000 permutations of the protein labels; features below it perform no better than chance. Features that vary together across the compared proteins ($|r|$, Cramér's V or correlation ratio $\geq 0.8$) are listed under the highest-ranked one and counted as one observation.

The source code of the pipeline and web application is available at https://github.com/TUNNELING-GROUP/LPMOcat and archived at https://doi.org/10.5281/zenodo.23698332; each pipeline step, its parameters and output files are documented at https://lpmo-cat.polsl.pl/docs/.

### Catalogue content

Table S1. Number of LPMOs per family, represented by experimental structures or predicted models.

| Family | Experimental | Predicted | Total |
|---|---|---|---|
| AA9 | 24 | 55 | 79 |
| AA10 | 27 | 25 | 52 |
| AA11 | 3 | 1 | 4 |
| AA13 | 1 | 10 | 11 |
| AA14 | 1 | 2 | 3 |
| AA15 | 1 | 4 | 5 |
| AA16 | 1 | 1 | 2 |
| AA17 | 3 | 2 | 5 |

| Family | Experimental | Predicted | Total |
|---|---|---|---|
| **All** | **61** | **100** | **161** |

Table S2. Taxonomic origin of the LPMOs in the catalogue.

| Taxon | Proteins | By family |
|---|---|---|
| Fungi · Ascomycota | 78 | AA9 60 · AA11 4 · AA13 11 · AA14 1 · AA16 2 |
| Fungi · Basidiomycota | 21 | AA9 18 · AA10 1 · AA14 2 |
| Oomycetes | 6 | AA15 1 · AA17 5 |
| Animals · Insecta | 4 | AA15 4 |
| Animals · Mollusca | 1 | AA10 1 |
| Plants · Streptophyta | 1 | AA10 1 |
| Bacteria · Actinomycetota | 20 | AA10 20 |
| Bacteria · Pseudomonadota | 16 | AA10 16 |
| Bacteria · Bacillota | 10 | AA10 10 |
| Viruses · Poxviridae | 3 | AA10 3 |
| Metagenomes | 1 | AA9 1 |

Table S3. Substrates of the LPMOs, as reported in papers or derived from EC numbers or protein names.

| Substrate | Reported in papers | From the EC numbers or the name | Total | By family |
|---|---|---|---|---|
| Cellulose | 57 | 49 | 106 | AA9 79 · AA10 20 · AA14 3 · AA15 2 · AA16 2 |
| Chitin | 17 | 31 | 48 | AA10 39 · AA11 4 · AA15 5 |
| Xyloglucan | 25 | 0 | 25 | AA9 24 · AA14 1 |
| Cello-oligosaccharides | 16 | 0 | 16 | AA9 15 · AA14 1 |
| Glucomannan | 12 | 0 | 12 | AA9 12 |
| Mixed-linkage glucan | 11 | 0 | 11 | AA9 11 |
| Starch | 5 | 6 | 11 | AA13 11 |
| Xylan | 11 | 0 | 11 | AA9 8 · AA14 3 |
| Plant biomass | 7 | 0 | 7 | AA9 7 |
| Pectin | 3 | 2 | 5 | AA17 5 |
| Cellulose derivatives | 4 | 0 | 4 | AA9 4 |
| Xyloglucan oligosaccharides | 3 | 0 | 3 | AA9 2 · AA14 1 |
| Mannan | 1 | 1 | 2 | AA9 1 · AA10 1 |
| Xylo-oligosaccharides | 2 | 0 | 2 | AA9 1 · AA14 1 |
| Chito-oligosaccharides | 1 | 0 | 1 | AA11 1 |

Table S4. Regioselectivity of the LPMOs, as reported in papers or derived from EC numbers.

| Regioselectivity | Reported in papers | From the EC numbers | Total | By family |
|---|---|---|---|---|
| C1 | 32 | 31 | 63 | AA9 27 · AA10 17 · AA11 4 · AA13 11 · AA14 2 · AA16 2 |
| C4 | 21 | 1 | 22 | AA9 19 · AA17 3 |
| C1/C4 | 29 | 45 | 74 | AA9 33 · AA10 35 · AA14 1 · AA15 5 |
| Unknown | 0 | 2 | 2 | AA17 2 |

Table S5. Sequence motifs following the N-terminal histidine of the histidine brace.

| Motif | Proteins | By family | Residues |
|---|---|---|---|
| H-G-X4-P-X2-R | 64 | AA10 48 · AA13 11 · AA15 4 · AA16 1 | H-G-Y-L 10 · H-G-Y-I 9 · H-G-Y-V 8 · H-G-S-V 6 · H-G-F-V 4 · H-G-S-M 4 · H-G-Y-M 3 · H-G-A-M 2 · H-G-R-L 2 · H-G-V-A 2 · H-G-W-V 2 · H-G-A-V 1 · H-G-F-I 1 · H-G-F-L 1 · H-G-G-L 1 · H-G-G-M 1 · H-G-H-L 1 · H-G-L-I 1 · H-G-R-V 1 · H-G-S-I 1 · H-G-V-T 1 · H-G-W-I 1 · H-G-Y-T 1 |
| H-G | 34 | AA9 30 · AA10 4 | H-G-Y-V 14 · H-G-F-V 6 · H-G-H-V 5 · H-G-A-V 2 · H-G-G-V 2 · H-G-H-I 1 · H-G-I-V 1 · H-G-Q-L 1 · H-G-Q-V 1 · H-G-W-I 1 |
| H-G-X4-P | 5 | AA17 5 | H-G-Y-I 3 · H-G-Y-M 1 · H-G-Y-V 1 |
| Other | 57 | AA9 48 · AA11 4 · AA14 3 · AA15 1 · AA16 1 | H-T-I-F 8 · H-Y-T-F 7 · H-A-I-F 5 · H-A-T-F 5 · H-A-T-V 3 · H-T-T-V 3 · H-Y-I-F 3 · H-M-K-M 2 · H-T-L-V 2 · H-T-R-V 2 · H-Y-T-L 2 · H-A-A-F 1 · H-A-Q-L 1 · H-A-T-W 1 · H-A-V-V 1 · H-I-A-F 1 · H-M-E-M 1 · H-M-M-M 1 · H-Q-I-H 1 · H-S-I-F 1 · H-S-T-F 1 · H-T-R-M 1 · H-V-V-A 1 · H-Y-F-F 1 · H-Y-T-W 1 · H-Y-V-F 1 |
| Not determined | 1 | AA9 1 | |

Table S6. Sequence motifs following the internal histidine of the histidine brace.

| Motif | Proteins | By family | Residues |
|---|---|---|---|
| H-X-G-P | 75 | AA9 70 · AA17 5 | H-H-G-P 17 · H-K-G-P 17 · H-P-G-P 13 · H-Y-G-P 7 · H-A-G-P 5 · H-L-G-P 3 · H-Q-G-P 3 · H-V-G-P 3 · H-I-G-P 2 · H-M-G-P 2 · H-N-G-P 1 · H-S-G-P 1 · H-W-G-P 1 |
| H-X-G | 32 | AA9 2 · AA10 9 · AA11 4 · AA13 11 · AA15 5 · AA16 1 | H-G-G-M 11 · H-G-G-G 4 · H-P-G-T 3 · H-P-G-D 2 · H-P-G-Y 2 · H-I-G-F 1 · H-K-G-T 1 · H-L-G-F 1 · H-L-G-R 1 · H-M-G-Y 1 · H-P-G-S 1 · H-P-G-W 1 · H-R-G-Y 1 · H-T-G-T 1 · H-W-G-Y 1 |
| H-X2-G-P | 5 | AA9 5 | H-A-Y-G-P 1 · H-E-Q-G-P 1 · H-P-Y-G-P 1 · H-S-Q-G-P 1 · H-T-I-G-P 1 |
| Other | 48 | AA9 1 · AA10 43 · AA14 3 · AA16 1 | H-G-A-D 7 · H-E-P-S 3 · H-K-T-S 3 · H-N-T-S 3 · H-A-T-S 2 · H-A-T-T 2 · H-F-D-D 2 · H-F-S-D 2 · H-N-P-S 2 · H-S-T-S 2 · H-S-T-T 2 · H-T-T-G 2 · H-V-T-K 2 · H-A-T-A 1 · H-A-T-K 1 · H-A-T-R 1 · H-H-P-G 1 · H-I-T-K 1 · H-K-D-N 1 · H-K-T-K 1 · H-K-T-R 1 · H-K-Y-R 1 · H-L-T-S 1 · H-R-T-A 1 · H-S-T-K 1 · H-S-T-N 1 · H-T-N-N 1 |
| Not determined | 1 | AA9 1 | |

Table S7. Second-sphere hydrogen-bonding motifs around the copper site.

| Motif | Proteins | By family |
|---|---|---|
| H-Q-Y | 85 | AA9 76 · AA10 7 · AA14 2 |
| R-E-F | 21 | AA10 21 |
| E-N-F | 19 | AA10 19 |
| Q-Q-Y | 16 | AA13 11 · AA17 5 |
| E-F | 4 | AA15 4 |
| N-E-Y | 4 | AA11 4 |
| E-A-F | 2 | AA10 2 |
| E-K-F | 2 | AA10 2 |
| Q-Y | 2 | AA9 1 · AA16 1 |
| E-Y | 1 | AA15 1 |
| H-E-Y | 1 | AA9 1 |
| N-F | 1 | AA10 1 |
| N-Q-Y | 1 | AA16 1 |
| N-Y | 1 | AA14 1 |
| Not determined | 1 | AA9 1 |

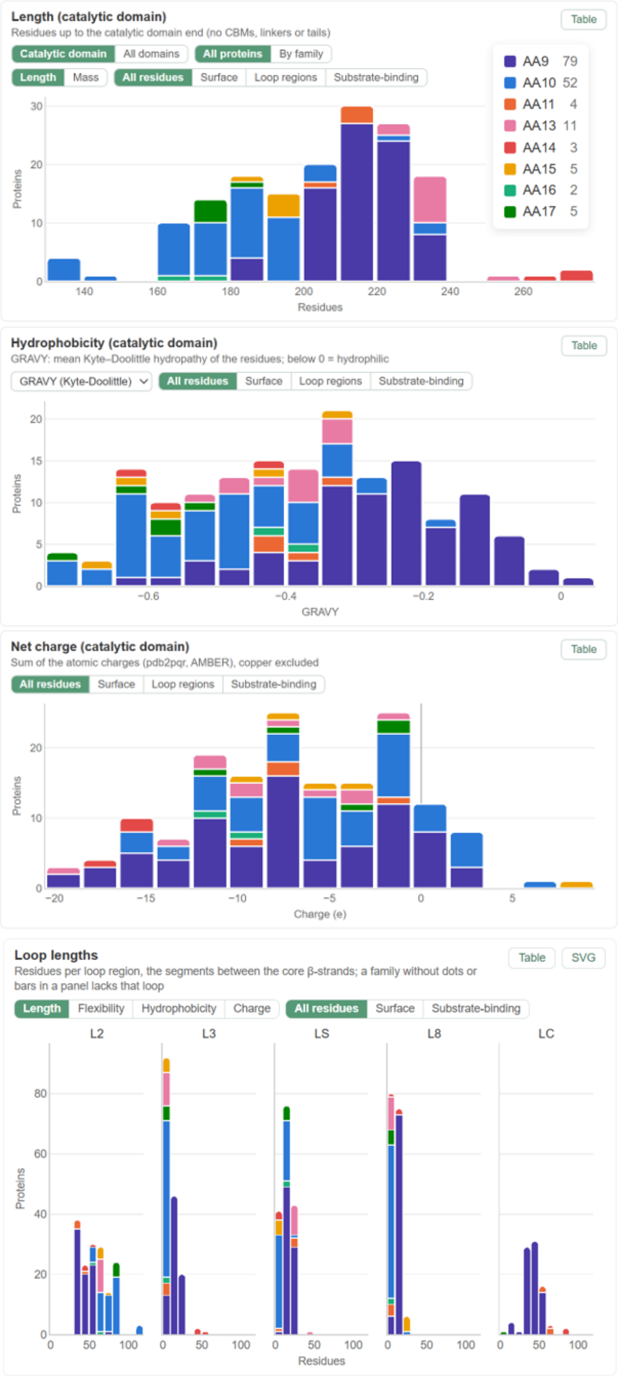

Figure S1. Properties of the catalytic domains across the catalogue. A) Length. B) Hydrophobicity (GRAVY, Kyte–Doolittle scale). C) Net charge (PDB2PQR, AMBER, copper excluded). D) Lengths of the loop regions L2, L3, LS, L8 and LC, defined as the segments between core β-strands; a family absent from a panel lacks that loop. Bars are coloured by family (AA9 purple, AA10 blue, AA11 orange, AA13 pink, AA14 red, AA15 yellow, AA16 teal, AA17 green).

| 39 | 1 | 30 | 24 | 15 |
|---|---|---|---|---|
| Proteins<br>of 161 in the catalogue | LPMO families<br>AA10 | Source species | Experimental structures<br>PDB | Predicted models<br>AlphaFold DB 14 · CAZyme3D 1 |

**Proteins**

Untick a protein to hide it in the 3D viewer. Click a column header to sort; hover it for its definition. Dotted underlined values show details on hover.

Show All ▾ Showing all 39 entries Columns ▾ Download table (.csv) Substrate evidence (.csv) **Download comparison (.zip)**

| | Name | Clade | CD length | Total length | CD mass (kDa) | Total mass (kDa) | CD net charge (e) | CD surface charge (e) | CD pI | CD hydrophobicity (GRAVY, Kyte–Doolittle) | Disulfide bonds | N-glyc. sequons | 3D structure | UniProt | Organism | Taxonomy |
|---|---|---|---|---|---|---|---|---|---|---|---|---|---|---|---|---|
| ☑ | SmAA10 | AA10 | 168 | 200 | 18.8 | 22.3 | 7.0 | 7.0 | 9.22 | -0.71 | 2 | 0 | AlphaFold DB Q59930 | Q59930 | Serratia marcescens | Bacteria +5 |
| ☑ | BpAA10A | AA10 | 196 | 196 | 22.1 | 22.1 | -6.0 | -8.0 | 5.60 | -0.64 | 0 | 0 | PDB 3UAM | Q3JY22 +1 | Burkholderia pseudomallei | Bacteria +5 |
| ☑ | LmAA10 | AA10 | 164 | 166 | 17.7 | 17.9 | -1.0 | 0.0 | 5.66 | -0.34 | 0 | 0 | PDB 5L2V | A0A0H3GIZ6 | Listeria monocytogenes | Bacteria +5 |
| ☑ | MaAA10 | AA10 | 194 | 194 | 21.6 | 21.6 | -2.0 | 1.0 | 6.19 | -0.51 | 2 | 0 | PDB 5OPF | UniParc UPI0001BF166D | Micromonospora aurantiaca | Bacteria +5 |
| ☑ | EfAA10A; EF0362 +2 | AA10 | 165 | 166 | 18.2 | 18.3 | 2.0 | 3.0 | 8.14 | -0.50 | 0 | 2 | PDB 4ALS +6 | Q838S1 +1 | Enterococcus faecalis | Bacteria +5 |
| ☑ | SliAA10E; SliLPMO10E +1 | AA10 | 172 | 172 | 18.4 | 18.4 | 2.0 | 5.0 | 8.42 | -0.45 | 2 | 2 | PDB 5FTZ | Q9RDB8 +2 | Streptomyces lividans | Bacteria +5 |
| ☑ | BtAA10A; Lpmo10A +1 | AA10 | 169 | 169 | 18.6 | 18.6 | 1.0 | 3.0 | 7.94 | -0.50 | 1 | 0 | PDB 5WSZ | A0A0C5K362 +2 | Bacillus thuringiensis | Bacteria +5 |

Figure S2. Summary table on the comparison page, shown for the 39 chitin-active AA10 LPMOs selected in Figure 1C. Tiles above the table summarise the selection (number of proteins, families and source species, experimental structures and predicted models). Each row lists the annotations and computed properties of one protein; columns can be shown or hidden, and the table scrolls horizontally when they do not fit the window. Superscript numbers link substrates, regioselectivity and EC numbers to the papers that report them, and “+n” expands further accessions or structure entries. Rows can be sorted and unticked to hide a protein from the 3D viewer and the plots. The table, the substrate evidence (including substrates tested without activity) and the whole comparison can be downloaded.

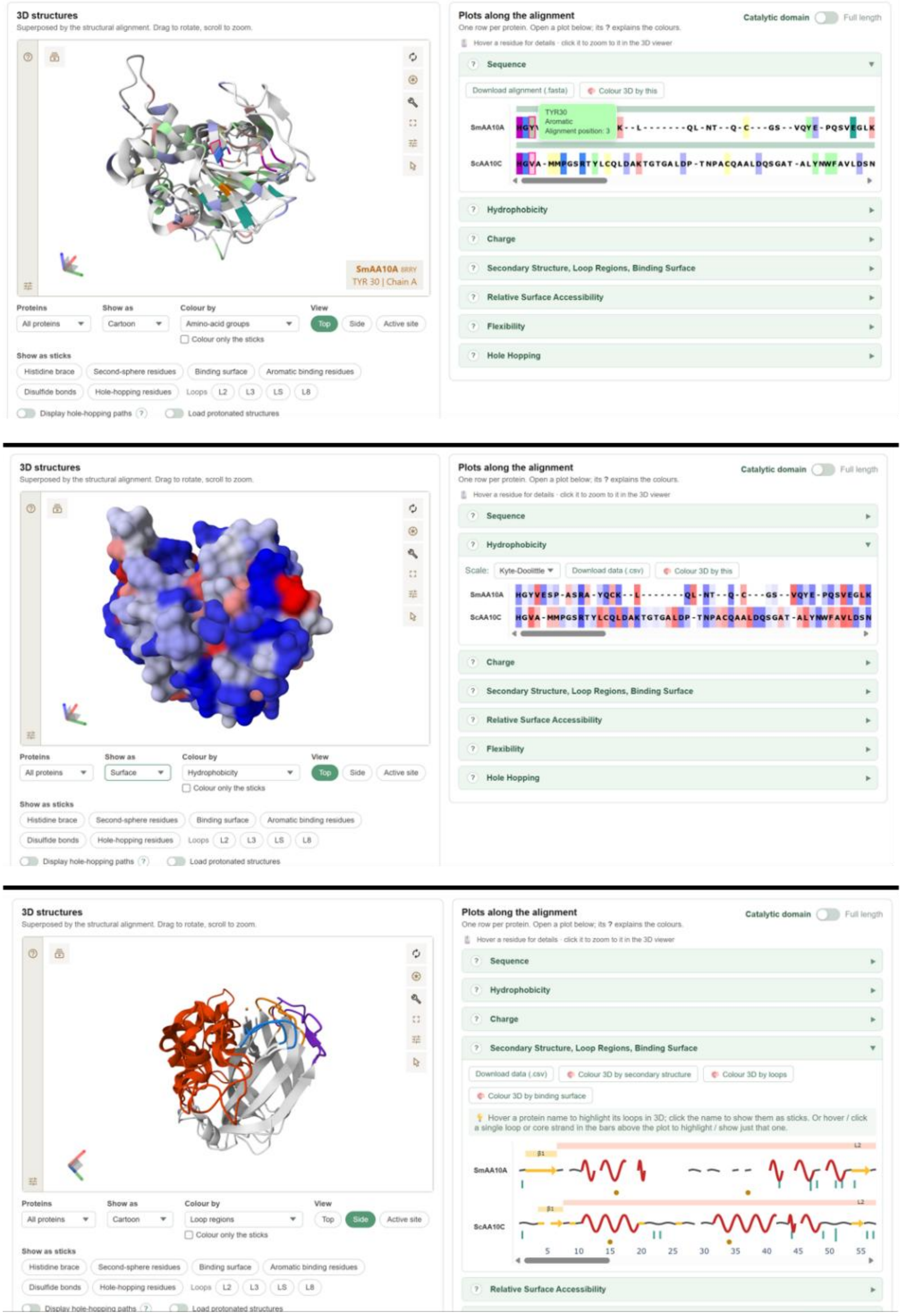

Figure S3. The 3D viewer and the plots along the alignment, shown for SmAA10A and ScAA10C. All structures are superposed in one frame, and each plot row is one protein aligned column by column. A) Structures coloured by amino-acid group. Pointing at a residue in the sequence plot (here Tyr30 of SmAA10A, an aromatic residue at alignment position 3) highlights it in the structure. B) Surface coloured by hydrophobicity (Kyte–Doolittle), with the matching hydrophobicity plot. C) Side view coloured by loop region (L2, L3, LS, L8, LC), with the secondary-structure track showing the core strands, loops and binding-surface residues.

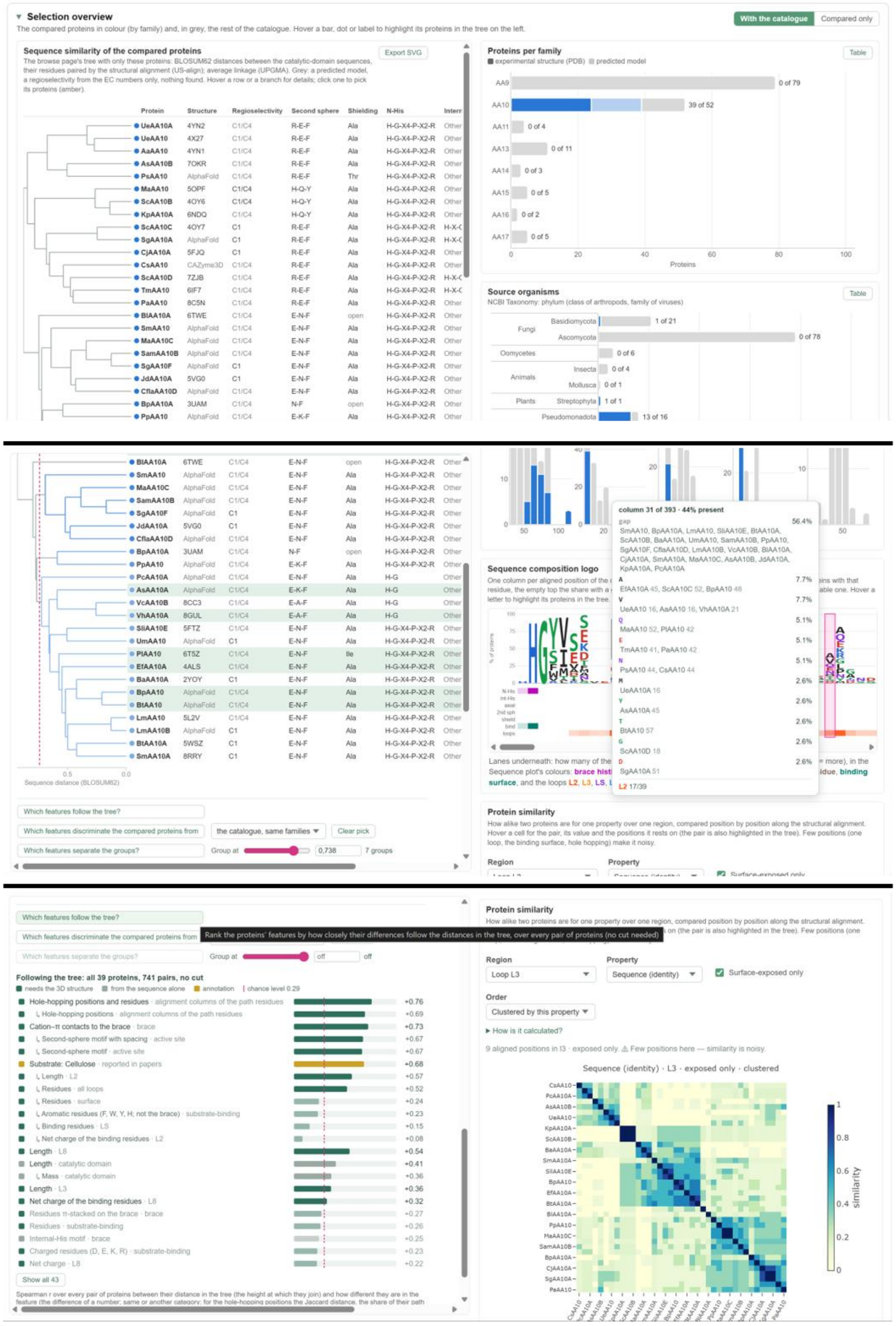

Figure S4. Selection overview for the 39 chitin-active AA10 LPMOs. A) The similarity tree pruned to the compared proteins, annotated with structure source, regioselectivity, second-sphere motif, shielding residue and brace-histidine motifs, next to the overview charts. The compared proteins are coloured by family and the rest of the catalogue is shown in grey. B) The tree cut into groups at a chosen distance (here 0.738, giving 7 groups), and the sequence logo of the selection with lanes marking the brace, second-sphere residues, binding surface and loops. Pointing at a column lists the proteins carrying each residue. C) Features ranked by how closely they follow the tree (dashed line, permutation-based threshold), and the pairwise similarity of the proteins for a chosen region and property (here sequence identity of the surface-exposed residues of loop L3, clustered by similarity).

Table S8. Features used to compare selected proteins with a reference set and with the similarity tree.

| No. | Feature | Definition | Unit | Basis |
|---|---|---|---|---|
| | **Catalytic domain** | | | |
| 1 | Length | Residues in the catalytic domain | residues | Seq |
| 2 | Mass | Molecular mass of the catalytic domain | kDa | Seq |
| 3 | Net charge | Sum of PDB2PQR (AMBER) atomic charges | e | Seq |
| 4 | Hydrophobicity | Kyte–Doolittle (GRAVY) scale | – | Seq |
| 5 | Isoelectric point | Computed with Biopython ProtParam | pH units | Seq |
| 6 | N-glycosylation sequons | Count of N-X-S/T motifs (X ≠ Pro); potential sites only | sites | Seq |
| | **Surface, loops and substrate-binding residues** | | | |
| 7 | Surface residues | Residues with relative SASA ≥ 25 % | residues | 3D |
| 8 | Surface net charge | Net charge of the surface residues | e | 3D |
| 9 | Surface hydrophobicity | Kyte–Doolittle (GRAVY) of the surface residues | – | 3D |
| 10 | Loop residues | Residues in all five loops together | residues | 3D |
| 11 | Loop net charge | Net charge of all loop residues | e | 3D |
| 12 | Loop hydrophobicity | Kyte–Doolittle (GRAVY) of all loop residues | – | 3D |
| 13 | Substrate-binding residues | Residues on the substrate-binding surface | residues | 3D |
| 14 | Substrate-binding net charge | Net charge of the substrate-binding residues | e | 3D |
| 15 | Substrate-binding hydrophobicity | Kyte–Doolittle (GRAVY) of the substrate-binding residues | – | 3D |
| 16 | Loop flexibility | Mean CABS-flex RMSF over all loop residues | Å | 3D |
| 17 | Flexibility of binding loop residues | Mean CABS-flex RMSF over the loops' substrate-binding residues | Å | 3D |
| | **Individual loops (each for L2, L3, LS, L8 and LC)** | | | |
| 18–22 | Loop length | Residues in the loop | residues | 3D |

| | | | | |
|---|---|---|---|---|
| 23–27 | Loop flexibility | Mean CABS-flex RMSF of the loop | Å | 3D |
| 28–32 | Loop net charge | Net charge of the loop | e | 3D |
| 33–37 | Loop hydrophobicity | GRAVY of the loop | – | 3D |
| 38–42 | Binding residues in the loop | Loop residues on the substrate-binding surface | residues | 3D |
| 43–47 | Net charge of binding residues in the loop | Net charge of those residues | e | 3D |
| | **Composition of the substrate-binding surface** | | | |
| 48 | Aromatic residues | Phe, Trp, Tyr and His among the substrate-binding residues, the two brace histidines excluded | residues | 3D |
| 49 | Amide residues | Asn, Gln among the substrate-binding residues | residues | 3D |
| 50 | Charged residues | Asp, Glu, Lys, Arg among the substrate-binding residues | residues | 3D |
| | Fold | | | |
| 51 | Disulfide bonds | Bonds with both Cys in the catalytic domain (S–S ≤ 3.0 Å) | bonds | 3D |
| 52 | Disulfide pattern | Regions (core strands, loops) joined by each bond, e.g. L2–β9 | category | 3D |
| 53 | β2 hairpin | Presence of the hairpin strand β2 after β1 | yes / no | 3D |
| | **Active site** | | | |
| 54 | Second-sphere motif | Hydrogen-bonding motif around the axial aromatic, e.g. H-Q-Y | category | 3D |
| 55 | Second-sphere motif with spacing | The same motif with residue spacing, e.g. H-X8-Q-X-Y | category | 3D |
| 56 | Shielding residue | Residue over the copper on its solvent-facing side; "open" if none | category | 3D |
| 57 | N-His motif | Sequence motif after the N-terminal brace histidine, e.g. H-G-X4-P-X2-R | category | Seq |

| | | | | |
|---|---|---|---|---|
| 58 | Internal-His motif | Sequence motif after the internal brace histidine, e.g. H-X-G-P | category | Seq |
| 59 | Brace hydrogen bonds | Residues whose side chain H-bonds a brace histidine (≤ 3.5 Å) | residues | 3D |
| 60 | Brace π-stacking | Residues π-stacked on a brace histidine | residues | 3D |
| 61 | Brace cation–π contacts | Residues in cation–π contact with a brace histidine | residues | 3D |
| 62 | N-His methylated | N-terminal His modelled as N-methylated His (HIC) in the PDB entry; no value for models | yes / no | 3D |
| | **Hole hopping** | | | |
| 63 | Hole-hopping positions | Alignment columns of the residues on all hole-hopping paths (EHPath); proteins compared by Jaccard distance | set of columns | 3D |
| 64 | Hole-hopping positions and residues | As 63, with a column shared only when both proteins have the same residue there | set of columns | 3D |
| | **Annotation** | | | |
| 65 | Carbohydrate-binding module | CBM families annotated in UniProt; "none" if absent | category | Ann |
| 66 | Regioselectivity | C1, C4 or C1/C4; from papers, otherwise from the EC number | category | Ann |
| 67 | Taxon | Taxonomic group and phylum (NCBI Taxonomy) | category | Ann |
| 68 | Structure source | Experimental structure or predicted model [a] | category | Ann |
| 69 | Cellulose | Activity reported in papers [b] | yes / no | Ann |
| 70 | Xyloglucan | Activity reported in papers [b] | yes / no | Ann |
| 71 | Chitin | Activity reported in papers [b] | yes / no | Ann |
| 72 | Cello-oligosaccharides | Activity reported in papers [b] | yes / no | Ann |
| 73 | Glucomannan | Activity reported in papers [b] | yes / no | Ann |
| 74 | Mixed-linkage glucan | Activity reported in papers [b] | yes / no | Ann |
| 75 | Xylan | Activity reported in papers [b] | yes / no | Ann |

Basis: Seq, computed from the sequence alone; 3D, needs the structure; Ann, from annotation.

All measures refer to the catalytic domain. Surface exposure is relative SASA ≥ 25 %. RMSF comes from CABS-flex simulations. A feature that cannot be computed for a protein (for example, no binding plane) is left without a value, not set to zero. Features that vary together across the proteins analysed (|Spearman r|, Cramér's V or correlation ratio ≥ 0.8) are shown under the highest-ranked one and counted as one observation.

[a] Not a biological property. It is included because it can create a grouping of its own.

[b] One feature for each substrate on which activity has been reported in papers for at least eight proteins in the catalogue. Proteins with no reported substrate have no value. "No" means not reported, which does not necessarily mean inactive.

### References

Abramson J, Adler J, Dunger J *et al.* Accurate structure prediction of biomolecular interactions with AlphaFold 3. *Nature* 2024;**630**(8016):493–500. https://doi.org/10.1038/s41586-024-07487-w.

Batka AE, Thomas WC, Tudorica DA *et al.* Second-Sphere Histidine Catalytic Function in a Fungal Polysaccharide Monooxygenase. *Biochemistry* 2024;**63**(23):3136–46. https://doi.org/10.1021/acs.biochem.4c00527.

Blum M, Andreeva A, Florentino LC *et al.* InterPro: the protein sequence classification resource in 2025. *Nucleic Acids Res* 2025;**53**(D1):D444–56. https://doi.org/10.1093/nar/gkae1082.

Bostock M, Ogievetsky V, Heer J. D$^3$ Data-Driven Documents. *IEEE Trans Vis Comput Graph* 2011;**17**(12):2301–9. https://doi.org/10.1109/TVCG.2011.185.

Burley SK, Bhikadiya C, Bi C *et al.* RCSB Protein Data Bank (RCSB.org): delivery of experimentally-determined PDB structures alongside one million computed structure models of proteins from artificial intelligence/machine learning. *Nucleic Acids Res* 2023;**51**(D1):D488–508. https://doi.org/10.1093/nar/gkac1077.

Cock PJA, Antao T, Chang JT *et al.* Biopython: freely available Python tools for computational molecular biology and bioinformatics. *Bioinformatics* 2009;**25**(11):1422–3. https://doi.org/10.1093/bioinformatics/btp163.

Dana JM, Gutmanas A, Tyagi N *et al.* SIFTS: updated Structure Integration with Function, Taxonomy and Sequences resource allows 40-fold increase in coverage of structure-based annotations for proteins. *Nucleic Acids Res* 2019;**47**(D1):D482–9. https://doi.org/10.1093/nar/gky1114.

Dolinsky TJ, Nielsen JE, McCammon JA *et al.* PDB2PQR: an automated pipeline for the setup of Poisson–Boltzmann electrostatics calculations. *Nucleic Acids Res* 2004;**32**(suppl_2):W665–7. https://doi.org/10.1093/nar/gkh381.

Drula E, Garron ML, Dogan S *et al.* The carbohydrate-active enzyme database: Functions and literature. *Nucleic Acids Res* 2022;**50**(D1):D571–7. https://doi.org/10.1093/nar/gkab1045.

Frandsen KEH, Lo Leggio L. Lytic polysaccharide monooxygenases: A crystallographer's view on a new class of biomass-degrading enzymes. *IUCrJ* 2016;**3**:448–67. https://doi.org/10.1107/S2052252516014147.

Hall KR, Joseph C, Ayuso-Fernández I *et al.* A Conserved Second Sphere Residue Tunes Copper Site Reactivity in Lytic Polysaccharide Monooxygenases. *J Am Chem Soc* 2023;**145**(34):18888–903. https://doi.org/10.1021/jacs.3c05342.

Hall KR, Mollatt M, Forsberg Z *et al.* Impact of the Copper Second Coordination Sphere on Catalytic Performance and Substrate Specificity of a Bacterial Lytic Polysaccharide Monooxygenase. *ACS Omega* 2024;**9**(21):23040–52. https://doi.org/10.1021/acsomega.4c02666.

Hauenstein J, Jeske L, Jäde A *et al.* BRENDA in 2026: a Global Core Biodata Resource for functional enzyme and metabolic data within the DSMZ Digital Diversity. *Nucleic Acids Res* 2026;**54**(D1):D527–34. https://doi.org/10.1093/nar/gkaf1113.

Hekkelman ML, Salmoral DÁ, Perrakis A *et al.* DSSP 4: FAIR annotation of protein secondary structure. *Protein Sci* 2025;**34**(8):e70208. https://doi.org/10.1002/pro.70208.

Hemsworth GR, Davies GJ, Walton PH. Recent insights into copper-containing lytic polysaccharide mono-oxygenases. *Curr Opin Struct Biol* 2013;**23**(5):660–8. https://doi.org/10.1016/j.sbi.2013.05.006.

Jumper J, Evans R, Pritzel A *et al.* Highly accurate protein structure prediction with AlphaFold. *Nature* 2021;**596**(7873):583–9. https://doi.org/10.1038/s41586-021-03819-2.

Kabsch W, Sander C. Dictionary of protein secondary structure: Pattern recognition of hydrogen-bonded and geometrical features. *Biopolymers* 1983;**22**(12):2577–637. https://doi.org/10.1002/bip.360221211.

Kurcinski M, Oleniecki T, Ciemny MP *et al.* CABS-flex standalone: a simulation environment for fast modeling of protein flexibility. *Bioinformatics* 2019;**35**(4):694–5. https://doi.org/10.1093/bioinformatics/bty685.

Laurent CVFP, Sun P, Scheiblbrandner S *et al.* Influence of Lytic Polysaccharide Monooxygenase Active Site Segments on Activity and Affinity. *Int J Mol Sci* 2019;**20**(24):6219. https://doi.org/10.3390/ijms20246219.

Levasseur A, Drula E, Lombard V *et al.* Expansion of the enzymatic repertoire of the CAZy database to integrate auxiliary redox enzymes. *Biotechnol Biofuels* 2013;**6**(1):41. https://doi.org/10.1186/1754-6834-6-41.

McDonald AG, Boyce S, Tipton KF. ExplorEnz: the primary source of the IUBMB enzyme list. *Nucleic Acids Res* 2009;**37**(Database):D593–7. https://doi.org/10.1093/nar/gkn582.

Plotly Technologies Inc. Collaborative data science. Montréal, QC, 2015. https://plot.ly.

Šali A, Blundell TL. Comparative Protein Modelling by Satisfaction of Spatial Restraints. *J Mol Biol* 1993;**234**(3):779–815. https://doi.org/10.1006/jmbi.1993.1626.

Sayler RI, Thomas WC, Rose AJ *et al.* Electron transfer in polysaccharide monooxygenase catalysis. *Proc Natl Acad Sci U S A* 2025;**122**(1). https://doi.org/10.1073/pnas.2411229121.

Schoch CL, Ciufo S, Domrachev M *et al.* NCBI Taxonomy: a comprehensive update on curation, resources and tools. *Database* 2020;**2020**:baaa062. https://doi.org/10.1093/database/baaa062.

Sehnal D, Bittrich S, Deshpande M *et al.* Mol* Viewer: modern web app for 3D visualization and analysis of large biomolecular structures. *Nucleic Acids Res* 2021;**49**(W1):W431–7. https://doi.org/10.1093/nar/gkab314.

Shrake A, Rupley JA. Environment and exposure to solvent of protein atoms. Lysozyme and insulin. *J Mol Biol* 1973;**79**(2):351–71. https://doi.org/10.1016/0022-2836(73)90011-9.

Siva Shanmugam NR, Yin Y. CAZyme3D: A Database of 3D Structures for Carbohydrate-active Enzymes. *J Mol Biol* 2025;**437**(15):169001. https://doi.org/10.1016/j.jmb.2025.169001.

Sokal RR, Michener CD. A statistical method for evaluating systematic relationships. *Univ Kans Sci Bull* 1958;**38**:1409–38.

Span EA, Marletta MA. The framework of polysaccharide monooxygenase structure and chemistry. *Curr Opin Struct Biol* 2015;**35**:93–9. https://doi.org/10.1016/j.sbi.2015.10.002.

Span EA, Suess DLM, Deller MC *et al.* The Role of the Secondary Coordination Sphere in a Fungal Polysaccharide Monooxygenase. *ACS Chem Biol* 2017;**12**(4):1095–103. https://doi.org/10.1021/acschembio.7b00016.

Teo RD, Wang R, Smithwick ER *et al.* Mapping hole hopping escape routes in proteins. *Proc Natl Acad Sci U S A* 2019;**116**(32):15811–6. https://doi.org/10.1073/pnas.1906394116.

Teufel F, Almagro Armenteros JJ, Johansen AR *et al.* SignalP 6.0 predicts all five types of signal peptides using protein language models. *Nat Biotechnol* 2022;**40**(7):1023–5. https://doi.org/10.1038/s41587-021-01156-3.

The UniProt Consortium. UniProt: the Universal Protein Knowledgebase in 2025. *Nucleic Acids Res* 2025;**53**(D1):D609–17. https://doi.org/10.1093/nar/gkae1010.

Tien MZ, Meyer AG, Sydykova DK *et al.* Maximum Allowed Solvent Accessibilites of Residues in Proteins. *PLoS ONE* 2013;**8**(11):e80635. https://doi.org/10.1371/journal.pone.0080635.

Varadi M, Bertoni D, Magana P *et al.* AlphaFold Protein Structure Database in 2024: providing structure coverage for over 214 million protein sequences. *Nucleic Acids Res* 2024;**52**(D1):D368–75. https://doi.org/10.1093/nar/gkad1011.

Zhang C, Shine M, Pyle AM *et al.* US-align: universal structure alignments of proteins, nucleic acids, and macromolecular complexes. *Nat Methods* 2022;**19**(9):1109–15. https://doi.org/10.1038/s41592-022-01585-1.

Zhang S, Krieger JM, Zhang Y *et al. ProDy* 2.0: increased scale and scope after 10 years of protein dynamics modelling with Python. *Bioinformatics* 2021;**37**(20):3657–9. https://doi.org/10.1093/bioinformatics/btab187.

Zhou X, Zhu H. Current understanding of substrate specificity and regioselectivity of LPMOs. *Bioresour Bioprocess* 2020;**7**(1). https://doi.org/10.1186/s40643-020-0300-6.
